# Single Nucleus Profiling Reveals Rewired Tumor-Immune-Stromal Cell Communication in Plasmacytoid Urothelial Carcinoma

**DOI:** 10.64898/2026.11.09.765457

**Authors:** Kathryn H. Gessner, Siyao Liu, Mi Zhou, Jeffrey S. Damrauer, KyuTae Hwang, David L. Corcoran, Brian C. Miller, Jonathan S. Serody, Hiroaki Murano, Minako Furusho, Kenichi Okuda, Wanda K. O’Neal, Hung-Jui Tan, Marc A. Bjurlin, Mary E. Westerman, Matthew E. Nielsen, Zachary S. Feuer, Angela B. Smith, Sara E. Wobker, Matthew I. Milowsky, William Y. Kim

## Abstract

Plasmacytoid urothelial carcinoma (PUC) is an aggressive histologic subtype of bladder cancer associated with worse patient outcomes and therapeutic response compared to conventional urothelial carcinoma (CUC). While recent work has demonstrated metastatic lymph nodes from PUC tumors have an inflamed phenotype, how the tumor microenvironment (TME) and crosstalk between tumor, stromal, and immune cells in primary PUC bladder tumors differs from those in CUC remains poorly understood. The impact of these differences on response to immunotherapy is also unknown. To understand TME differences between CUC and PUC and their impact on cell signaling, we performed single nucleus RNA sequencing on PUC and CUC tumors, as well as Xenium spatial transcriptomics on a mixed CUC / PUC tumor. We identified increased signaling between PUC-enriched, CDH1-low urothelial cells, immunosuppressive lipid-associated macrophages, and SFRP2+/CTHRC1+ fibroblasts in PUC tumors compared to CUC. Spatial transcriptomics identified increased macrophage infiltration in PUC histology and colocalization of PUC tumor cells, immunosuppressive macrophages, and SFRP2+ fibroblasts. Taken together, this study delineates PUC architecture and demonstrates interplay CDH1-low tumor cells, EMT, and immunosuppressive macrophages, identifying possible therapeutic vulnerabilities in PUC.

## INTRODUCTION

Plasmacytoid urothelial carcinoma (PUC) is a rare and aggressive subtype of bladder cancer with up to 5% of urothelial carcinoma tumors containing a component of PUC histology.^1–3^ Patients with PUC typically present with higher stage disease at diagnosis and have a worse prognosis compared to conventional urothelial carcinoma (CUC).^4–6^ Additionally, PUC demonstrates poor response to neoadjuvant cisplatin-based chemotherapy. PUC response to enfortumab vedotin plus the immune checkpoint inhibitor pembrolizumab (EVP), now first-line therapy for peri-operative muscle-invasive and advanced bladder cancer, remains uncertain, highlighting the need for therapeutic development^7,8^. Improved understanding of the underlying biology of PUC will allow for informed therapeutic selection or development of targeted therapies to improve outcomes for patients.

The characteristic alteration in PUC is *CDH1* mutation in up to 84% of tumors and reduced or loss of E-cadherin protein expression by immunohistochemistry in up to 100% of PUC.^9–14^ The high rate of CDH1 mutation frequency in PUC contrasts with CUC, in which 0% of the TCGA CUC cohort demonstrated truncating CDH1 mutations.^9^ E-cadherin is a transmembrane protein involved in cell-cell adhesion whose loss is tightly linked to epithelial-to-mesenchymal transition (EMT).^15^ PUC tumors express mesenchymal markers, including Snail and vimentin, suggesting induction of EMT.^12^ Apart from *CDH1* mutations, the mutational landscape of PUC is similar to CUC, with just minor differences in the rates of p53 and RB1 mutations (RB1: 55% vs 17%, p<0.001; TP53: 76% vs 48%, p=0.009).^16^ This similarity in mutational landscape highlights the importance of the canonical loss of E-cadherin protein in PUC.

In other solid tumors, upregulation of EMT impacts the composition of cell types present in the tumor microenvironment (TME), such as tumor associated macrophages (TAMs),^17,18^ regulatory T-cells^19^ and cancer-associated fibroblasts (CAFs)^20^. However, prior characterization of the TME in PUC is limited. A recent report from a rapid autopsy program demonstrated metastatic lymph nodes from PUC tumors are immune-inflamed, whereas the TME of primary PUC tumors remains less well characterized.^21^ With the establishment of EVP as standard of care for perioperative therapy in muscle-invasive bladder cancer and first-line therapy for advanced and metastatic disease, understanding the immune microenvironment of PUC primary tumors is critical to understanding the potential efficacy of these therapies as well as the potential development of new approaches.

We hypothesized that loss of E-cadherin and upregulation of EMT in PUC tumors promotes alterations in immune contexture to create therapeutic vulnerabilities. In this study, we characterize changes in the TME in PUC tumors compared to CUC tumors and define the molecular pathways involved. Specifically, CDH1-low PUC tumor cells demonstrate increased interactions with immunosuppressive CTHRC1+SFRP2+ fibroblasts and lipid-associated SPP1+/TREM2+ macrophages compared to CUC tumors. By understanding the changes in the TME and crosstalk between tumor and immune cells at high resolution, we aim to identify potential therapeutic vulnerabilities that lead to the development of novel treatments with improved efficacy and outcomes for PUC, a particularly lethal histologic subtype of bladder cancer.

## METHODS

### Tissue staining and selection

Patient tissue samples were obtained under IRB protocol #23-1112, which was approved by the University of North Carolina Institutional Review Board. A waiver of informed consent was approved by the IRB. Given the small number of samples, sex was not considered as a biological variable in this study. Patient tissue samples were distributed by the Pathology Services Core. H&E stains were performed using the autostainer XL from Leica Biosystems and chromogenic immunohistochemistry (IHC) was performed on freshly obtained paraffin-embedded sections. E-cadherin IHC was carried out using the Leica Bond Rx Autostainer system. Slides were deparaffinized in Bond Dewax solution (AR9222) and hydrated in Bond Wash solution (AR9590). Heat induced antigen retrieval was performed at 100°C with Bond-Epitope Retrieval solution 2 pH-9.0 (AR9640) for 20 minutes. After pretreatment, slides were blocked and then incubated with ready-to-use E-Cadherin antibody (Leica Biosystems, PA0387) for 2 hours followed by ready-to-use Novolink Polymer Detection Kit (Leica Biosystems, RE7260-CE) with Novolink Polymer plus Post Primary secondaries. Antibody detection with 3,3’-diaminobenzidine (DAB) was performed using the Bond Intense R detection system (DS9263). Stained slides were dehydrated and coverslipped with Cytoseal 60 (Epredia, 83104). Stained slides were digitally imaged in the Aperio AT2 (Leica Microsystems) using 20x objective and interpreted by a genitourinary pathologist using ImageScope (Leica Microsystems).

### Single nucleus RNA sequencing

Tumor areas from CUC and UC containing plasmacytoid histology were identified. Based on histology, samples were classified as CUC, pure PUC, or mixed PUC/CUC by a genitourinary pathologist (SEW). Pure PUC classification indicated that no conventional histology was observed in the tissue sections utilized for single nucleus RNA sequencing (snRNAseq), whereas mixed CUC/PUC contained both histologies in the tissue area. RNA was isolated from FFPE tissue sections using the Qiagen RNeasy FFPE kit following manufacturer’s instructions. DV200 was assessed using a High Sensitivity RNA ScreenTape assay and 4150 TapeStation instrument (Agilent). Nuclei were isolated from FFPE tissue sections, stored at - 80C, and thawed following 10x Genomics Demonstrated Protocol CG000632 Rev D with the pestle dissociation method. Aliquots were stained with acridine orange and propidium iodide and assessed for concentration and singleness using the LUNA-FX7 Dual Fluorescence Cell Counter (Logos Biosystems). The Chromium Fixed RNA Profiling Next-GEM Reagent Kit (10x PN-1000476) was then used to perform sample multiplexing, probe-based RNA capture, and library preparation following 10x Demonstrated Protocol CG000527 Rev E. Library quality was assessed using the High Sensitivity DNA 5000 ScreenTape assay (Agilent). Paired-end sequencing was performed on the Illumina NextSeq2000 by the Translational Genomics Laboratory at UNC.

### Computational analysis of single nuclear RNA sequencing

Raw data was processed and a Seurat object was generated as previously described.^22^ Cells with unique features counts < 200 or < 8000 or with > 5% mitochondrial counts were filtered out. The data were log-normalized and top 2000 variable features were identified. Cells from tumors were clustered individually and Uniform Manifold Approximation and Projection (UMAP) dimensional reduction was performed. Based on established markers of cell types, distinct clusters were classified as a general cell type. snRNAseq data from individual tumors were then integrated using Harmony integration in Seurat. Cells within the integrated dataset were then clustered using resolution 0.1 and UMAP dimensional reduction analysis was performed. Epithelial, myeloid, macrophage, T-cells, and fibroblast populations were further subset using the same process to identify subclusters of these cell types. Cell proportions were compared between histologies using the two-sided pairwise Wilcoxon rank-sum test, implemented with the R function state_compare_means() (method = “wilcox.text”). P-values from these comparisons were not adjusted for multiple comparisons.

Markers of each subpopulation were identified using the FindAllMarkers function in Seurat. Gene signatures were calculated using the addModule function from Seurat and UCell package in R. Differential analysis was performed using FindMarkers function in Seurat. GSEA pathway enrichment was performed using clusterProfiler package in R. Trajectory analysis was performed using monocle2 using DDR tree method on selected genes. Cellular communication analysis was performed using the CellChat package (v2.1.2) in R. Ligands and receptors that are significantly associated with cellular communication with PUC-enriched tumor cells to macrophages were annotated on a pathway level using EnrichR.^23^

### Single cell spatial transcriptomics

In situ spatial transcriptomic analysis was conducted on paraffin embedded samples. Pre-existing H&E images with selected areas of interest were reviewed to guide sectioning and ensure optimal tissue orientation and placement of regions of interest. After the specialized Xenium slides were equilibrated to room temperature, tissue samples were sectioned at 5um using RNAse free conditions. Two different samples were placed on each Xenium slide, with a total of 4 samples analyzed across 2 Xenium slides in a single run, one of which is utilized for this study. This assay was carried out using the manufacturer guidelines for the Xenium Analyzer (10x Genomics) with the Human Multi-Tissue and Cancer Panel plus UN92PC Xenium add on Custom Probe. Slides were loaded onto the Xenium Analyzer, where transcripts were detected through automated cycles of probe hybridization and imaging. Resulting images were visualized in Xenium Explorer, with punctate transcript signals displayed as color-coded markers across the tissue.

### Computational analysis of Xenium spatial transcriptomics

#### Quality control

Cell-by-feature matrices were obtained from the Xenium Analyzer (10X Genomics; onboard analysis) and analyzed in R (v4.3.2) using Seurat package (v5.1.0). Cells with fewer than 10 transcripts and genes with fewer than 10 total transcripts across all cells were removed Data was analyzed using Seurat R package. Expression values were log-normalized using the NormalizeData() function (method = “LogNormalize” scale factor = 10,000), and scaled using the ScaleData() function. Principal component (PC) analysis was performed on all 477 panel genes retained after filtering using the RunPCA() function, and the first 30 PCs were used to construct a UMAP embedding.

#### Clustering and cell type annotation

Clusters were identified using the FindNeighbors() and FindClusters(). Coarse and fine level cell labels were derived independently from three soures: the graph-based clustering produced by the Xenium Analyzer and visualized in Xenium Explorer (v4.1.0), Seurat clustering, and BANKSY (v1.2.0). A consensus label with then assigned to each cell type by manually reviewing the canonical marker expression of each cell type across the three sources. Macrophages were subset and re-clustered in Seurat (re-running PCA on the same subet, using 30PCs, resolution = 0.2), and the subclusters were visualized by UMAP. Differentially expressed genes between subclusters were identified using FindAllMarkers() function (Wilcoxon rank-sum test, min.pct=0.1, logFC threshold=0, Bonferroni-adjusted P < 0.05). The top differentially expressed genes in the PUC-specific macrophage and fibroblast cluster were used as gene signatures. These signatures were used to score the corresponding cell types in the snRNAseq dataset with UCell. UCell was utilized to assign a module score to the snRNAseq clusters for these cell types. RNA density plots and images demonstrating *in situ* transcript localization were generated using Xenium Explorer.

#### Cellular neighborhood analysis

Neighborhood analysis was performed using the hoodscanR package (v1.0.0) in R. The processed and annotated Seurat object was converted to a SpatialExperiment object, containing spatial coordinates (cell centroids). For each cell, the 100 nearest neighboring cells were identified with findNearCells(), which uses the Approximate Nearest Neighbor (ANN) algorithm implemented in the RANN package. Neighborhood probability distributions were computed with scanHoods() which uses the proximity-focused softmax algorithm, and probabilities were summarized by coarse- and fine-level cell type labels with mergeByGroup(). Neighborhood entropy and perplexity were calculated with calcMetrics() as measures of local cellular diversity. Pairwise colocalization between cell types was quantified as the Pearson correlation of their neighborhood probabilities across cells. Positive and negative values were interpreted as spatial co-occurrence and segregation, respectively. Cells were clustered by their neighborhood probability profiles with K-means visa clustByHood() for k = 2–10. Tissue maps and probability distribution plots were inspected for each clustering solution, and k=5 was selected for downstream interpretation based on biologically meaningful neighborhood spatial organization.

#### Nearest-neighbor distance analysis

To quantify the spatial proximity of urothelial cell populations to tumor microenvironment cell types, we computed distances for each plasmacytoid urothelial carcinoma (PUC) cell and each conventional urothelial cell (CUC). For each of these cells, we measured the Euclidean distance (µm) to the nearest cell of each target cell type. The target types were myeloid cells, fibroblasts (all fibroblasts and the Fibroblast_1 and Fibroblast_2_KI67 subclusters), endothelial cells, B cells, and T cells. Distances were calculated using the nndist() function from spatstat.geom (v3.5.0) in R. The resulting minimum distance distributions were compared between PUC and CUC populations using boxplots, and statistical significance was assessed with the Wilcoxon rank-sum test as implemented in ggpubr::stat_compare_means().

### Analysis of TCGA BLCA dataset

Based on the findings from the snRNAseq data, tumors from The Cancer Genome Atlas Bladder Cancer study^24^ were analyzed to evaluate the clinical outcomes associated with SFRP2 and CTHRC1 expression levels. RSEM normalized counts were utilized and the median expression of each gene across all tumor histologies was calculated. Tumors above the median expression level were classified as “high” expression, while tumors with expression level below the median value were classified as “low”. Statistical tests included Kruskal-Willis to assess the significance of expression level by tumor stage and Cox Hazard proportion analysis to assess the differences in overall survival.

### Data availability

Data will be made available in a public repository at time of publication.

## RESULTS

### Single nucleus RNA sequencing reveals PUC-enriched subclusters of epithelial cells with CDH1 loss

Evaluation of PUC tumor cell biology and the TME within PUC tumors compared to CUC tumors was performed through snRNAseq of tumor areas deemed histologically PUC-only (4), CUC-only (4), mixed PUC/CUC (4) (Figure 1). Cell clusters were identified and broad cell types were assigned using canonical markers for individual cell types (Figure 2A, Supplemental Figure 1). Within immune cells, the proportion of general cell types is not significantly different between CUC and PUC tumors (Figure 2B). To further interrogate gene expression differences in urothelial cells between CUC and PUC tumors, epithelial cells were subclustered and resolved into 6 urothelial cell subclusters (Figure 2C). Cluster 3 demonstrated low-quality metrics and was removed from further analyses (Supplemental Figure 2A). Of the remaining 5 clusters, Cluster 1 demonstrates a basal-like pattern of gene expression, higher CDH1 expression, and enrichment in CUC tumors (Figures 2D,E,F). Clusters 0 and 2 demonstrate enrichment in PUC and mixed PUC/CUC tumors, higher expression of luminal markers, and lower CDH1 expression (Figures 2D,E,F). Within clusters 0 and 2, PUC tumors have low levels of CDH1 expression, while CUC tumors have higher levels (Figure 2F). Additionally, clusters 0 and 2 demonstrate higher levels of MUC16, MUC4, and TM4SF1 (Figure 2F), consistent with a previous report demonstrating higher expression of these genes in urothelial variant histologic subtypes.^25^

**Figure 1.**
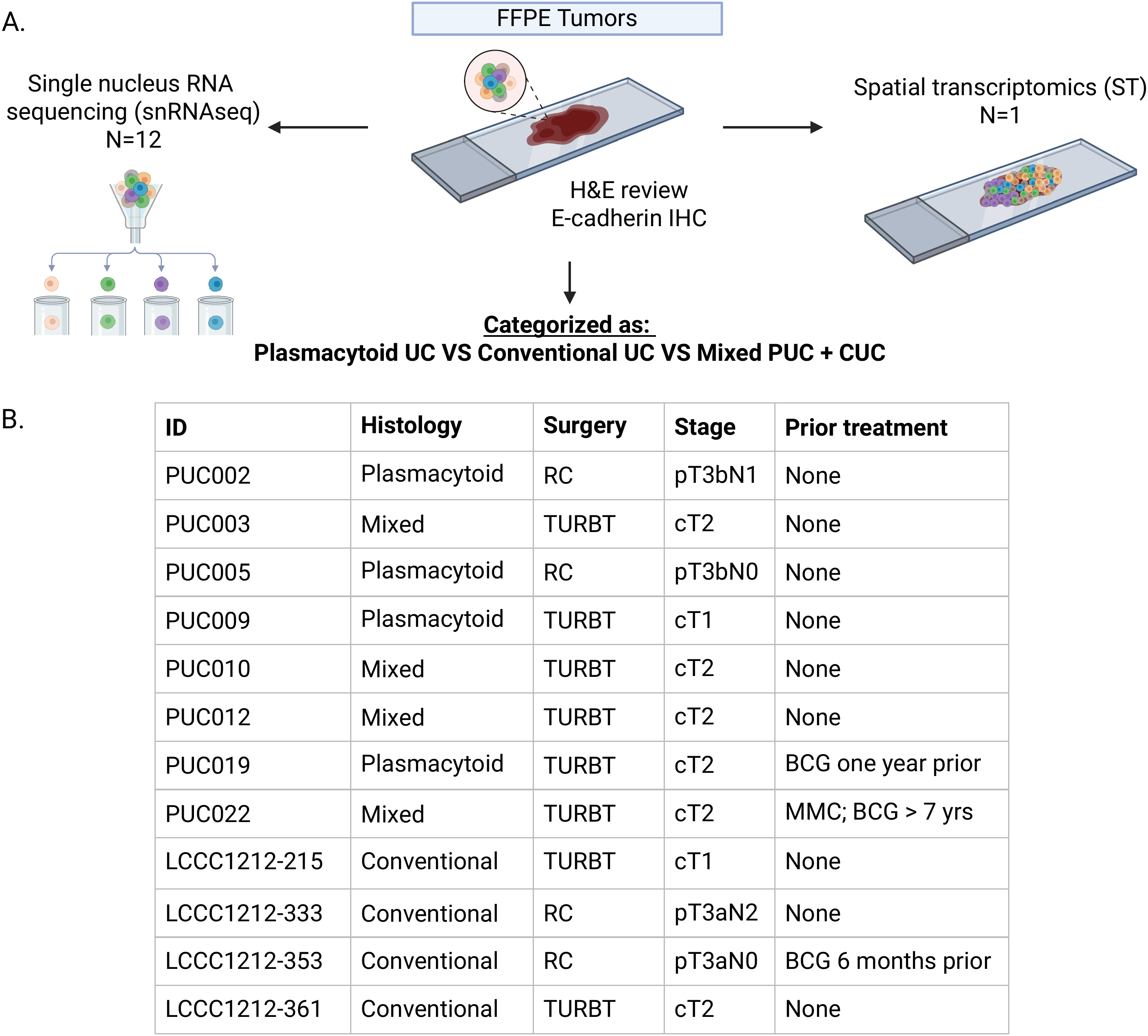
Study design and cohort overview. (A) Schematic of study workflow. (B) Characteristics of individual tumors included in the study, including histology of tumor tissue used in molecular assays, type of surgery, tumor stage (pathologic for cystectomy tumors, clinic for TURBT tumors), and previous treatment history.

**Figure 2.**
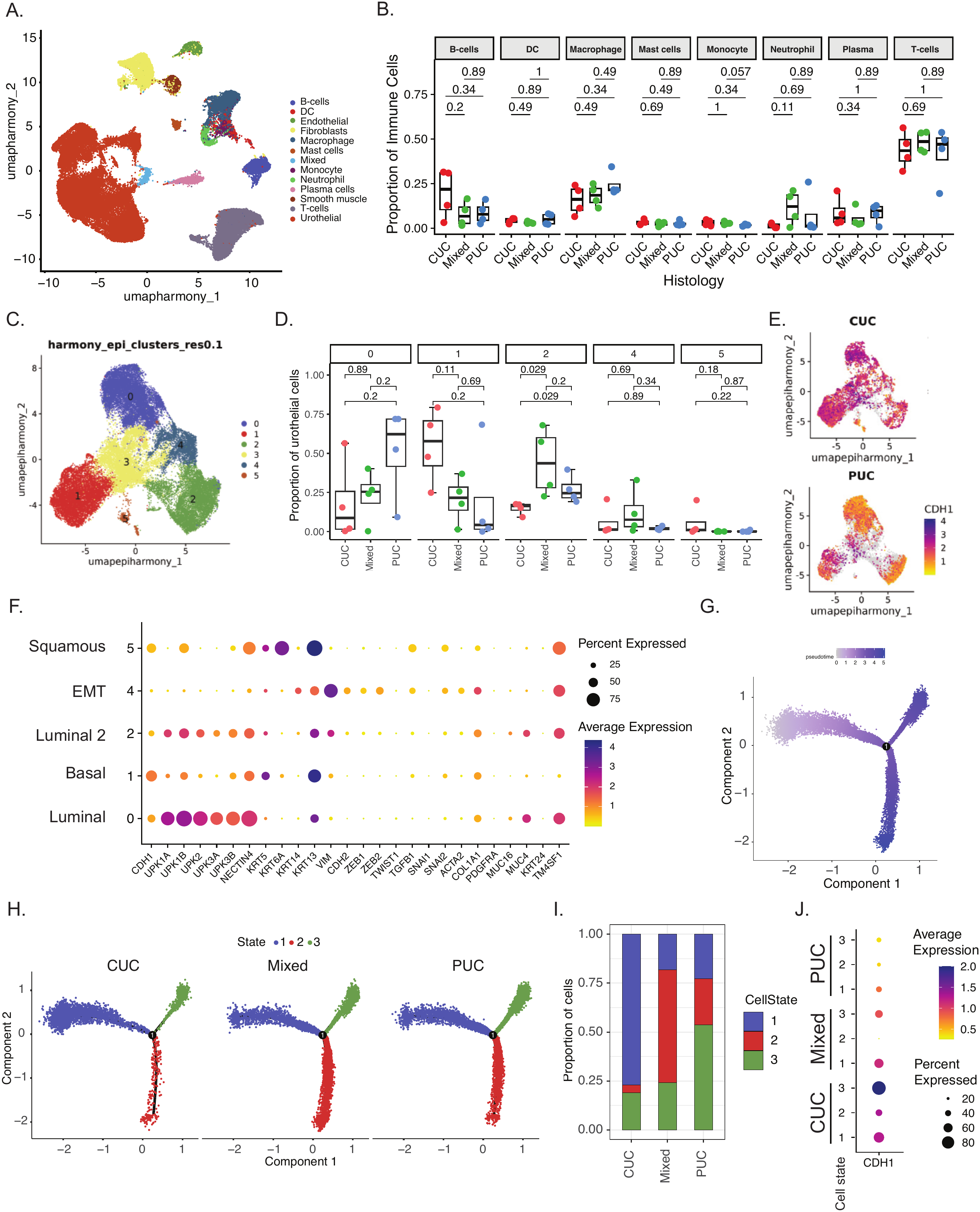
snRNAseq identifies CDH1-low urothelial cell populations enriched in PUC tumors with differentiated, luminal phenotype. (A) UMAP of general cell types across all tumor samples. (B) Boxplot showing proportion of immune cell types in PUC, CUC, and mixed tumors. Boxes indicate the 25^th^ to 75^th^ percentiles, the center line the median, and whiskers extend to 1.5 x IQR; points are individuals tumors. P-values by Wilcoxon rank-sum test. (C) UMAP of urothelial cell types across all tumor samples colored by urothelial subclusters. (D) Boxplot showing proportion of each urothelial subcluster of all urothelial cells, split by CUC (red), PUC (blue), and mixed (green) tumors. Boxes indicate the 25^th^ to 75^th^ percentiles, the center line the median, and whiskers extend to 1.5 x IQR; points are individuals tumors. P-values by Wilcoxon rank-sum test. (E) Feature plot showing CDH1 expression in urothelial cells, split by CUC tumors (top) and PUC tumors (bottom). (F) Dot plot of select genes stratified by urothelial subcluster. Size of the circle indicates percent of cells expressing each gene. Color of circle indicates average gene expression across cells of that subcluster. (G) Trajectory inference analysis of urothelial cell populations plotted by pseudotime. (H) Trajectory inference analysis of urothelial cell populations plotted by cell state of CUC (left), mixed (middle), and PUC (right) tumors. (I) Stacked bar plot comparing the proportion of urothelial cells classified as each CellState, stratified by tumor histology. (J) Trajectory inference analysis of urothelial cell populations colored by CDH1 expression level, split by tumor histology

To further characterize the differentiation pattern of the tumor epithelial cells in CUC vs PUC histology, trajectory inference analysis was performed. This analysis demonstrates a luminal differentiation pattern that progresses to node 1 and then splits into two populations (Figure 2G). When plotted by CellState, this progression shows increased luminal transcriptional phenotype when moving from CellState 1 to CellStates 2 and 3, with a distinct branchpoint between CellState 2 and 3 (Figure 2H). CUC tumors are enriched for CellState 1, while mixed and PUC tumors are enriched for CellStates 2 and 3, respectively (Figure 2H,I), consistent with prior reports that PUC have a luminal phenotype.^26,27^ Across tumor histologic subgroups, CellStates 2 and 3 demonstrates lower CDH1 expression (Figure 2J). To better define the differences between CellStates 2 and 3, we performed differential gene expression between these Cell States. Cell State 2 demonstrates higher expression of genes involved in EMT (HMGA2, AXL, VIM, WWTR1 (TAZ), FOSL1, HGF), cancer stem cell-like markers (CD44 and ANPEP), and ECM remodeling (ADAMTSL1, MMP14, PLAU), as well as MAGEA4, a target of a bispecific T cell engager in development (Supplemental Figure 2B).^28^ Gene Set Enrichment Analysis demonstrates higher expression of pathways involved in Epithelial to Mesenchymal Transition, inflammatory response, and interferon alpha and gamma response in Cell State 2 compared to Cell State 3 (Supplemental Figure 2C). On the contrary, Cell State 3 retains more epithelial cell identity, as evidenced by higher KRT20, UPK1A, UPK3A, CD24, GATA3, FOXA1, and CDH1 expression than Cell State 2 (Supplemental Figure 2). Cell State 3 also demonstrates expression of pathways involved in metabolic reprogramming and estrogen response (Supplemental Figure 2). The combined differential expression and trajectory analysis demonstrate urothelial populations enriched in PUC histology contain a more luminal transcriptional state than CUC histology and express higher levels of partial EMT activation, ECM remodeling, and lineage plasticity. Concurrent upregulation of these programs supports an invasive, treatment-resistant phenotypic state, consistent with plasmacytoid biology.

### Analysis of the tumor microenvironment and cellular communication identifies increased signaling between PUC-enriched urothelial clusters, macrophage populations, and fibroblast populations

To characterize the TME within PUC tumors, immune and fibroblast cell populations were further subset. Within the T-cell cluster, 6 distinct subpopulations were identified and assigned T-cell identities based on canonical T-cell markers (Supplemental Figure 3A,B). Proportions of T-cell subclusters are not significantly different between PUC, mixed, and CUC tumors (Supplemental Figure 3C). As the relationships between CD8+ T-cells, macrophages, and fibroblasts are associated with response to immunotherapy, we examined the ratio of CD8+ T-cells to these other main cell types. The ratio of TAMs to CD8+ T-cells is numerically increased in PUC tumors compared to CUC tumors (Supplemental Figure 4A). The ratio of CD8+ T-cells to fibroblasts is numerically decreased in PUC vs CUC tumors (Supplemental Figure 4B). Higher TAM/CD8+ T-cell and lower CD8+ T-cell/CAF ratios are associated with worse response to immunotherapy.^29,30^ Macrophage and fibroblast populations were further subset and classified as cellular subtypes based on canonical markers (Supplemental Figures 4C-D,5A-B). PUC tumors demonstrate a significantly higher proportion of APOE+/GPNMB+ macrophages than CUC tumors (Supplemental Figure 4E). GPNMB+ macrophages are an immunosuppressive subset of macrophages that co-localize with terminally exhausted CD8+ T-cells and attenuate CD8+ T-cell response.^31–33^ There are no significant differences in proportions of fibroblast subclusters in PUC compared to CUC tumors (Supplemental Figure 5C). The findings of a high TAM/CD8+ T-cell ratio and the increased proportion of APOE+/GPNMB+ macrophages in PUC tumors suggest a more immunosuppressive TME.

We next performed CellChat analysis to understand the differences in cellular communication between tumor epithelial cells and other cell types in PUC compared to CUC tumors. In PUC tumors compared to CUC tumors, urothelial cell populations enriched in PUC-tumors (urothelial c0 and c2) and CTHRC1+SFRP2+ fibroblasts (c0) demonstrate increased interactions with lipid-associated (LA), TREM2+SPP1+ macrophages and inflammatory macrophages (macrophage c2 and c6, respectively) in PUC tumors compared to CUC tumors (Figure 3A,B). LA and inflammatory macrophages also demonstrate increased signaling to PUC-enriched urothelial cell populations and themselves, suggesting bi-directional crosstalk between these cell populations (Figure 3A,B). CellChat analysis identifies ligand-receptor pairs involved in signaling between PUC-enriched urothelial cell populations to LA-TAMs (Figure 3C).

**Figure 3.**
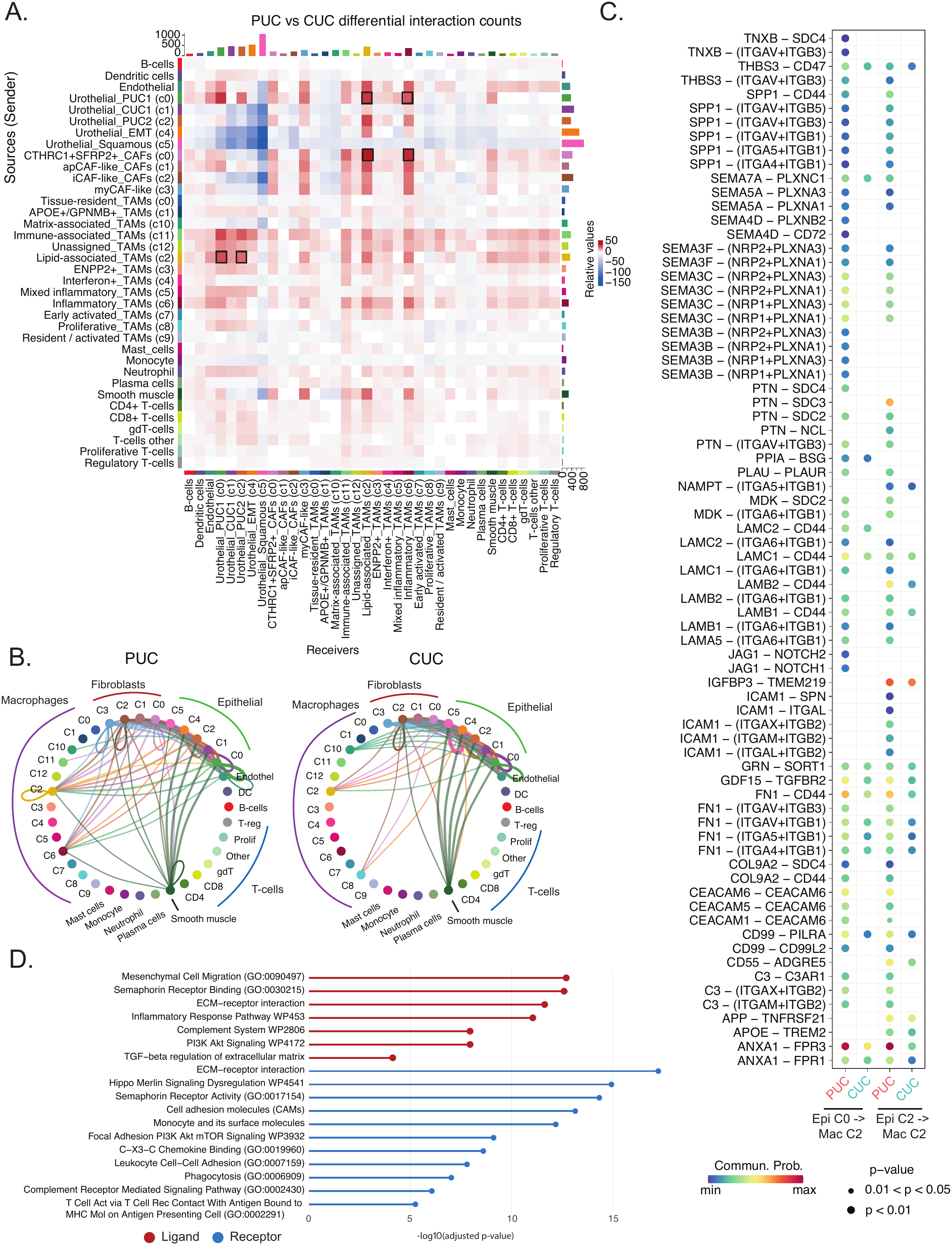
Cellular interaction analysis identifies increased signaling between plasmacytoid-enriched urothelial cell populations, lipid-associated and inflammatory macrophages, and SFRP2+/CTHRC1+ fibroblasts. (A) Heatmap of differential interaction counts between PUC vs CUC tumors. Sources of ligands shown on the y-axis and receiving cells expressing receptors are on the x-axis. Black boxes indicated interactions of interest. (B) Circle plot demonstrating interactions between cell types in PUC tumors (left) vs CUC tumors (right). Each node is a cell cluster. Arrows go from sending cell type to receiving cell type and the arrow width indicates the number of interactions. (C) Bubble plot of ligand-receptors pairs predicted as involved in signaling from urothelial clusters c0 and c2 to LA-TAMs in PUC vs CUC. Bubble size indicates p-value. Bubble color indicates communication probability. (D) Lollipop plot of molecular pathways identified as enriched by EnrichR in ligands (red) and receptors (blue) predicted as involved in signaling from urothelial clusters c0 and c2 to LA-TAMs in PUC vs CUC.

To identify the molecular pathways enriched in these interactions, ligands and receptors were analyzed using EnrichR. Pathways involved in signaling between PUC-enriched epithelial cells (c0 and c2) and LA macrophages include those associated with extracellular matrix signaling and adhesion, semaphorin signaling, complement and innate immunity, chemokine binding, and pathways associated with EMT (PI3K/Akt, Hippo, TGF-beta) (Figure 3D). Ligand-receptor pairs with significantly increased signaling between c0 fibroblasts to c2 and c6 macrophages, urothelial c0 and c2 to macrophage c6, and macrophage c2 to urothelial c0 and c2 were identified (Supplemental Figure 6). These data suggest that PUC-enriched epithelial cells have distinct interactions with immunosuppressive cell types in the TME compared to CUC-enriched epithelial cells. We hypothesize that these interactions promote the aggressive tumor phenotypes and therapeutic resistance seen in PUC.

### Single cell resolution spatial transcriptomics identifies plasmacytoid histology as a distinct transcriptomic entity

To further interrogate the spatial proximity of these cellular relationships, we performed spatial transcriptomics on a bladder tumor with mixed conventional and plasmacytoid histology using the 10X Genomics Xenium spatial transcriptomics platform, which allows for single cell resolution. We first characterized this tumor by H&E to identify the location of each histologic subtype and performed IHC for E-cadherin protein. This tumor demonstrates loss of E-cadherin protein expression by immunohistochemistry in the plasmacytoid histology components (Figure 4A). Graph-based dimensional reduction analysis identified broad clusters (Figure 4B) which were assigned into 11 coarse cell types (Figure 4C). The assigned cell types demonstrate the expected expression of canonical cell type markers (Supplemental Figure 7). When cell identities were projected onto the *in-situ* locations, areas of plasmacytoid and conventional urothelial carcinoma histology were identified as distinct transcriptomic clusters (Figure 4C).

**Figure 4.**
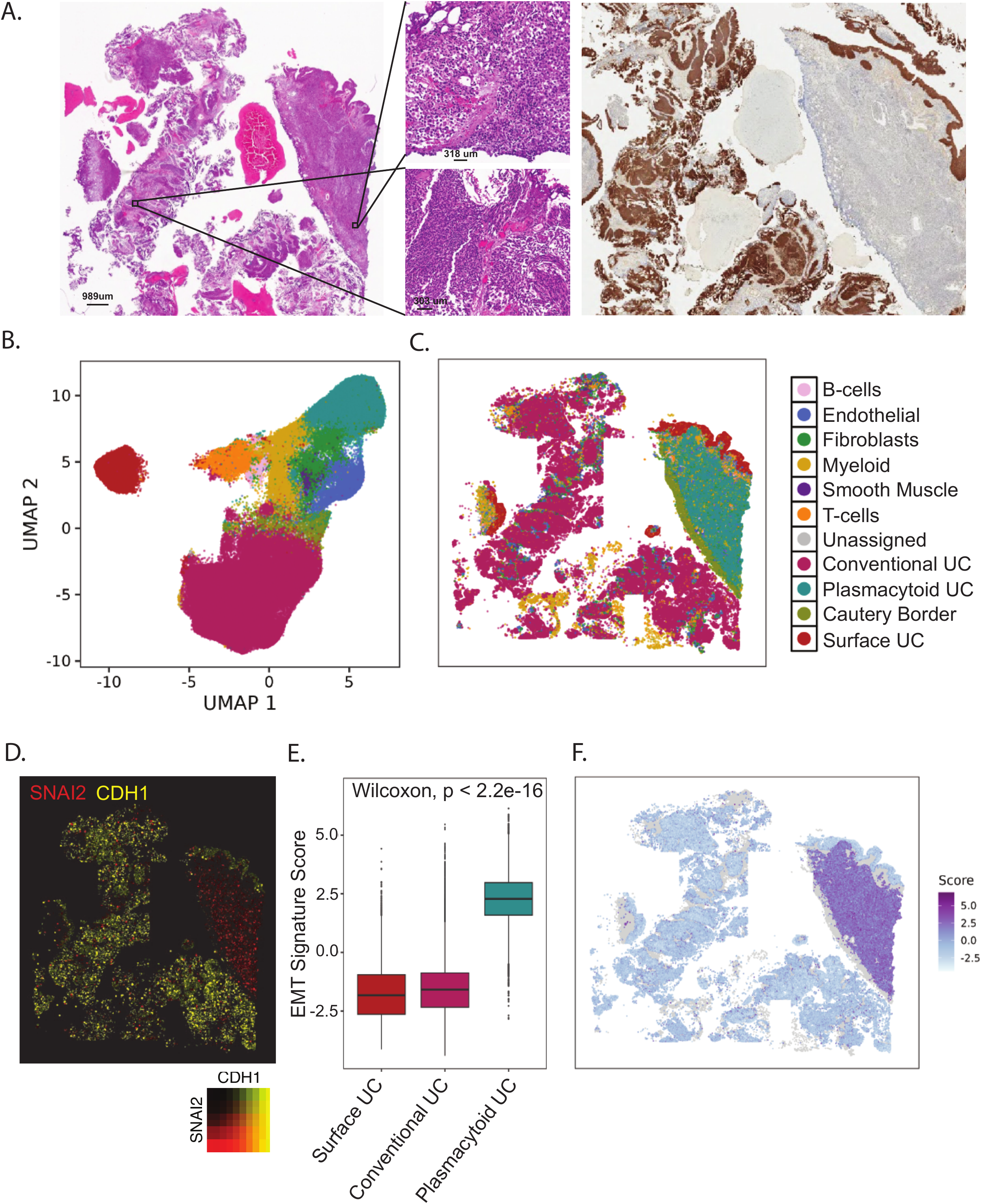
Xenium spatial transcriptomics on a mixed CUC / PUC tumor demonstrates loss of *CDH1* and increased expression of mesenchymal markers in the PUC histologic region. (A) *Left:* H&E image of PUC003 mixed histology tumor with PUC region on right (top inset) and CUC region on left (bottom inset). *Right*: E-cadherin immunohistochemistry staining. (B) UMAP of general cell types. (C) Spatial plot of general cell types. (D) Spatial feature plot of *CDH1* (yellow) and *SNAI2* (red) expression. (E) Boxplot demonstrating the expression of the EMT signature score published by Yauch *et al*, split by urothelial subcluster. P-value calculated using Wilcoxon rank-sum test. (F) Spatial feature plot of the EMT signature score (Yauch *et al*) across tissue image.

We then evaluated transcriptomic differences between plasmacytoid and conventional histology urothelial carcinoma. As expected, PUC histology demonstrates loss of *CDH1* expression and increased expression of the mesenchymal marker, SNAI2 (Figure 4D). The PUC cluster also demonstrates significantly increased expression of a previously published EMT signature score^34^ (Figures 4E,F, p < 2.2e-16), confirming upregulation of EMT in plasmacytoid histology.

### Neighborhood analysis identifies increased infiltration of myeloid cells, fibroblasts, and T-cells in plasmacytoid histologic region

To interrogate the snRNAseq findings, we next evaluated the spatial relationships between tumor cells and immune cells in plasmacytoid and conventional histology. Evaluating the *in-situ* image, plasmacytoid histology demonstrates a visually apparent increase in myeloid cells, T-cells, and fibroblasts (Figure 5A). Spatially resolved neighborhood analysis using HoodScanR identified 5 spatially distinct cellular niches: a CUC-neighborhood, PUC-neighborhood, stromal neighborhood, surface urothelium neighborhood, and neighborhood associated with likely cautery artifact (Figure 5B). The PUC-neighborhood demonstrates a higher probability of macrophages, T-cells, and fibroblasts being present within the spatially resolved neighborhood relative to the CUC-neighborhood (Figure 5C). This finding was orthogonally validated by calculating the distance between individual PUC and CUC cells and the closest neighboring cell of each immune or stromal subgroup. PUC cells showed a significantly shorter distance to T-cells, myeloid cells, fibroblasts, and B-cells compared to CUC cells (p < 2.2e-16, Figure 5D). Therefore, the spatial composition of PUC neighborhoods differs from CUC neighborhoods, with increased complexity and immune and stromal composition.

**Figure 5.**
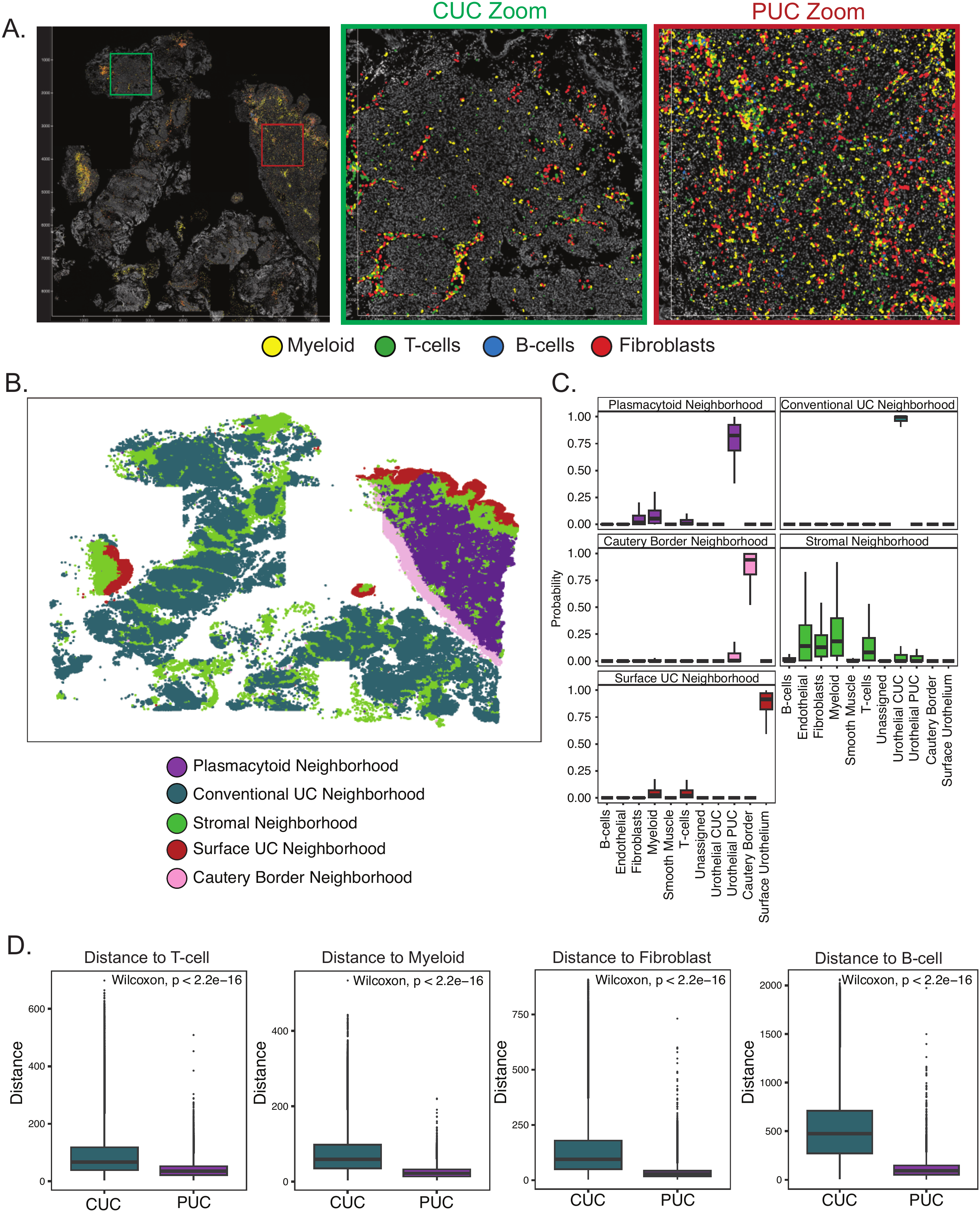
Plasmacytoid region of a mixed histology tumor demonstrates increased infiltration of myeloid cells and fibroblasts. (A) Spatial plot of myeloid cells (yellow), T-cells (green), B-cells (blue), and fibroblasts (red) with inset of CUC region (green outline, middle) and PUC region (red outline, right). (B) Spatial plot demonstrating locations of distinct spatial neighborhoods defined using unsupervised clustering in CellChat. (C) Box plots demonstrating the probability of each general cell type within the five spatial neighborhoods. Box plot color indicates neighborhood identity. (D) Box plots showing the distance from each individual CUC (teal) or PUC (purple) cell to other cell types (*left to right*: T-cell, myeloid, fibroblast, B-cell). Significance testing completed using Wilcoxon rank sum.

### Communicating cellular populations in PUC tumors are located in close spatial proximity

To further characterize the spatial proximity of immune and stromal cells within PUC histology, myeloid cells were subset and subclustering was performed. Six macrophage populations were identified in the Xenium data (Figure 6A) and Cluster 1 localized to the region of plasmacytoid histology (Figure 6B). When we applied a module score of the top differentially expressed genes in the Xenium macrophage c1 cluster (Figure 6B) to the snRNAseq macrophage clusters (Supplemental Figure 4C,D), snRNAseq cluster 2 matched the Xenium macrophage c1 population (Figure 6B), consistent with it demonstrating increased cellular communication with PUC-enriched tumor cells. Similarly, two transcriptomic clusters of fibroblasts were identified in the Xenium data (Figure 6D). Cluster c2 localizes to PUC histology and, when compared to CUC histology, demonstrates significantly higher expression of SFRP2 (average log2 fold-change = 4.58; adjusted p-value = 0), as well as ligands identified as involved in cellular communication with LA macrophages in the snRNAseq data: THY1 (1.74, 0), SEMA3C (1.49, 7.05e-301) and ITGB5 (1.06, 1.02e-283) (Figure 6E, Supplemental Figure 8).

**Figure 6.**
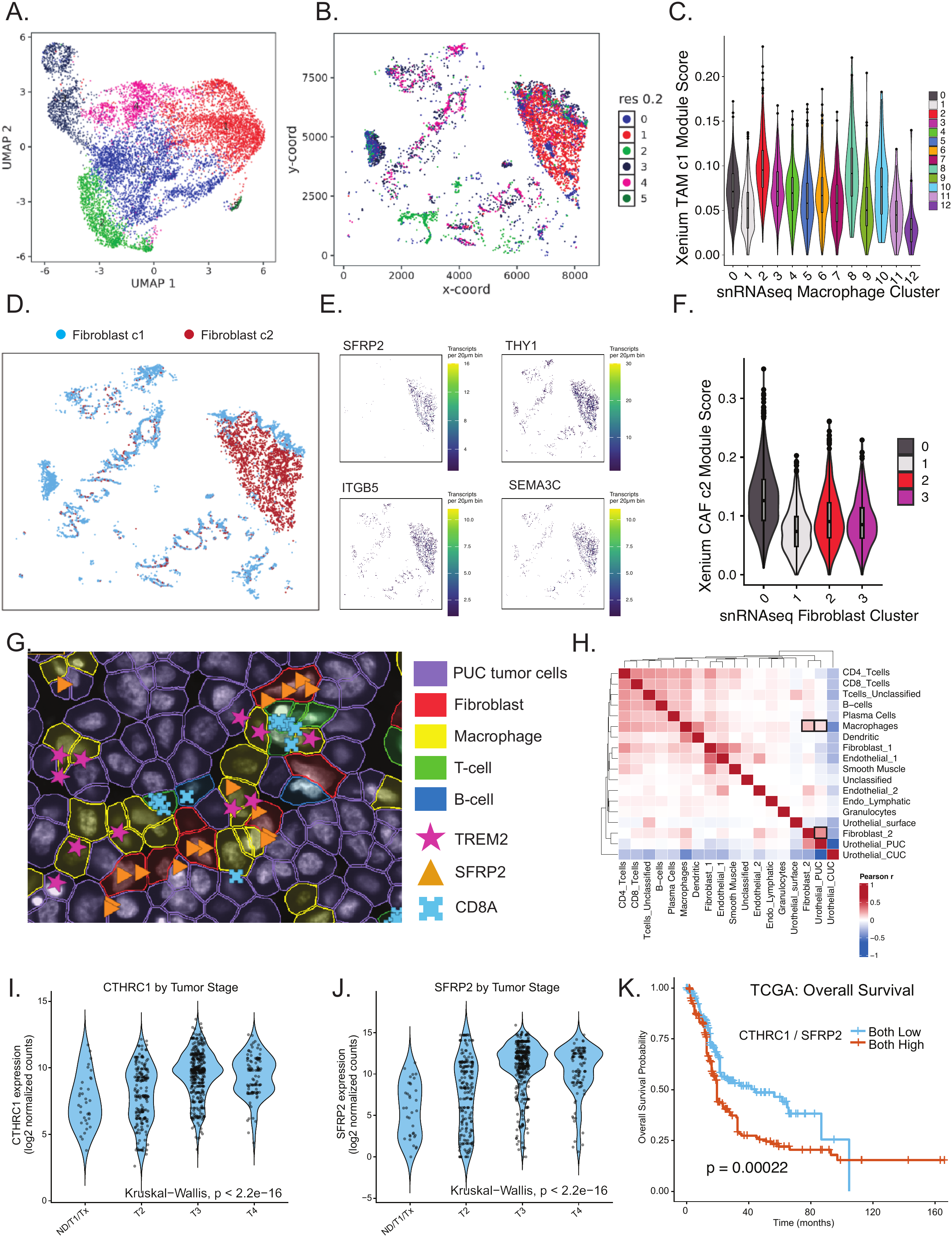
Lipid-associated (TREM2+/SPP1+/APOE+) macrophages, SFRP2+/CTHRC1+ fibroblasts, and CDH1-tumor cells are located in close spatial proximity within region of plasmacytoid histology. (A) UMAP of macrophage subclusters showing annotated identities in Xenium data. (B) Spatial plot of macrophage subclusters. (C) Violin plot depicting the module score of the c1 macrophage subcluster from Xenium data, stratified by the snRNAseq macrophage clusters. (D) Spatial plot of fibroblast subclusters in the Xenium data from the PUC003 tumor. (E) Spatial feature plots of the expression of genes relevant to PUC-associated fibroblasts in the snRNAseq data (SFRP2, THY1, ITGB5, SEMA3C). Cell color represents Q-score density. (F) Violin pot depicting the module score of the c2 fibroblast subcluster from Xenium data, stratified by the snRNAseq fibroblast clusters. (G) Xenium Explorer image showing a region of interest from the PUC histology region depicting individual cells. Cell color and shading indicate cell identity. Transcript identity and location for specific genes of interest is depicted (TREM2: pink star, SFRP2: orange triangle, CD8A: blue notched square). (H) Neighborhood enrichment analysis between cell types in spatial coordinates, calculated by Squidpy. Box color indicates the enrichment z-score. (I) Violin plot of CTHRC1 expression in tumors from the bladder cancer TCGA cohort, stratified by tumor stage. Y-axis depicts CTHRC1 log2 normalized counts. P-value is calculated using Kruskal-Willis test. (J) Violin plot of SFRP2 expression in tumors from the bladder cancer TCGA cohort, stratified by tumor stage. Y-axis depicts SFRP2 log2 normalized counts. P-value is calculated using Kruskal-Willis test. (K) Kaplan-Meier analysis of overall survival (OS) in the bladder cancer TCGA cohort, stratified by CTHRC1 and SFRP2 expression, using a median split and comparing tumors that have high expression of both genes versus those with low expression of both genes.

When a module score of this Xenium-based cluster was applied to the snRNAseq fibroblast clusters, snRNAseq fibroblast cluster 0 was most consistent with this fibroblast identity (Figure 6F). While the probe panel for Xenium did not include CTHRC1, expression of SFRP2 was specific for the CTHRC1+SFRP2+ fibroblast cluster in the snRNAseq dataset. Therefore, we utilized this specificity to identify the CTHRC1+SFRP2+ fibroblast population within the Xenium *in situ* data. A representative *in situ* image of the plasmacytoid region within the Xenium data demonstrates close spatial proximity of TREM2+ macrophages, SFRP2+ fibroblasts, and PUC tumor cells (Figure 6G), providing corroborating evidence that the interactions observed in the snRNAseq data reflect genuine biological relationships. HoodScanR colocalization heatmap demonstrates spatial colocalization of these cell types (Figure 6H).

To further interrogate the clinical relevance of the presence of CTHRC1+SFRP2+ cells, the association between gene expression and clinical features was evaluated within the TCGA BLCA dataset. TCGA was utilized because we lack a large genomic dataset with plasmacytoid urothelial carcinoma and clinical outcomes. CTHRC1 and SFRP2 expression are independently significantly increased in higher tumor stages (Kruskal-Willis, p < 2.2e-16) (Figures 6I, J, respectively). Patients with bladder cancer that demonstrates high expression of both CTHRC1 and SFPR2 have a significantly shorter overall survival than patients with tumors with low expression of both genes (Cox proportional hazard, p = 0.0002) (Figure 6K). While the TCGA is majority CUC, this association supports that CTHCR1+SFRP2+ fibroblasts are associated with poor clinical outcomes.

## DISCUSSION

This study provides novel insights into plasmacytoid tumor cell biology and the PUC TME. Consistent with prior reports, PUC tumor cells demonstrate a luminal phenotype and expression of MUC4 and TM4SF1, stem cell markers expressed at higher levels in histologic bladder cancer subtypes.^25–27^ PUC tumor cells also exhibit a partial EMT state and retain markers of epithelial cell origin. Within the PUC TME, our data reveals rewired tumor-stromal-immune cell communication with CDH1-low tumor cells showing stronger predicted signaling to immunosuppressive TAMs and CAFs than their CUC, CDH1-high counterparts.

The current study identifies two populations of PUC tumor cells with CDH1 loss which demonstrate varying levels of EMT activation and urothelial transcriptional phenotypes. While EMT has traditionally been treated as two binary states (epithelial vs mesenchymal), recent work has highlighted the plasticity of EMT, with cells existing along a continuous spectrum.^35^ Cell populations with varying levels of EMT have been previously reported in cancer tissue.^36^ Canonical, or full, EMT is traditionally due to loss of E-cadherin reportedly due to transcriptional repression, which activates mesenchymal markers, but also requires a decrease in other epithelial proteins, including cytokeratins.^36,37^ Partial EMT has been associated with persistence of epithelial lineage markers, low ZEB1 expression, epigenetic changes that stabilize the partial EMT state, and E-cadherin internalization.^36,38^ In PUC tumors, urothelial populations with different levels of EMT may be due to varying levels of *CDH1* RNA (Supplemental Figure 2), timing of E-cadherin loss, differences in signaling from surrounding stroma or immune cells, or differences in other cell-intrinsic signaling or mutations.^36,39^

PUC tumors harbor an immunosuppressive TME with increased cellular signaling between CTHRC1+SFRP2+ fibroblasts, SPP1+TREM2+ macrophages, and CDH1-low tumor cells. CTHRC1+ and SFRP2+ fibroblasts are immunosuppressive fibroblast populations associated with poor patient outcomes in other tumor types, consistent with our finding in the TCGA BLCA dataset.^40–42^ In pancreatic ductal adenocarcinoma, CTHRC1+GREM1+ fibroblasts and SPP1+APOE+ macrophages synergistically promote an immunosuppressive tumor microenvironment.^42^ Similarly, CellChat analysis in the current study inferred interactions among CTHRC1+SFRP2+ fibroblasts, SPP1+TREM2+ macrophages, and CDH1-low tumor cells, supporting the hypothesis that an analogous synergistic immunosuppressive circuit occurs in PUC tumors. Although these interactions are inferred from transcriptional data and require functional validation, they nominate a cooperative circuit promoting an immunosuppressive environment in PUC, which may contribute to its aggressive biology and poor patient outcomes.

While PUC has historically demonstrated worse response to chemotherapy than CUC, it remains uncertain whether PUC will respond to EVP. To date, small studies and reports demonstrate conflicting reports of response to EV monotherapy, immune checkpoint inhibitor monotherapy, and EVP.^3,8,16,43–46^ The current study provides biologic insights into PUC response to EVP. PUC-enriched tumor cells demonstrate increased expression of NECTIN4, the target of EV, suggesting potential on-target activity. However, snRNAseq cannot assess membranous NECTIN4 protein expression, which is more directly relevant for predicting EV response.^47^ On the contrary, PUC tumors also demonstrate a higher TAM/CD8+ T-cell ratio and lower CD8+ T-cell/CAF ratio, which are associated with worse response to immunotherapy.^29,30^ Additionally, the signaling among CDH1-tumor cells and immunosuppressive populations of macrophages and fibroblasts discussed above also predict poor response to immunotherapy.^41,42,48,49^ Together, these findings suggest that PUC harbors features of both potential response and resistance to EVP, necessitating further interrogation of EVP response in preclinical PUC models.

This work also provides insight into novel therapeutic targets or mechanisms that warrant further testing in PUC tumors. As described by Yang *et al.*, PUC tumors express high levels of TM4SF1, a surface protein that acts as a cell cycle and apoptosis regulator.^25^ TM4SF1-CAR T-cells have *in vitro* activity against bladder cancer cell lines, highlighting its potential as a therapeutic target.^25^ Furthermore, PUC tumor cells demonstrate activated TGF-beta signaling, NOTCH signaling, and activation of the TAP/TAZ pathway, all of which have inhibitors in development.^50–52^ In addition to tumor-cell specific targets, the study identifies possible therapeutic vulnerabilities in the PUC TME. As SFRP2+ fibroblasts are associated with poor response to combination anti-PD-1/anti-HER2 therapy, combination SFRP2 neutralizing antibody plus immunotherapy is being tested in a Phase 1b study.^41,53^ Similarly, treatment of an ovarian cancer model with an anti-TREM2 monoclonal antibody increases CD8+ T-cell infiltration and response to anti-PD-1 therapy, highlighting the potential role for combining TREM2+ inhibition with immunotherapy in PUC.^48^ Finally, TREM2+SPP1+ LA macrophages in PUC tumors also express PLA2G7, an immunosuppressive TAM phenotype with worse response to immune checkpoint blockade.^54,55^ Treatment of PLA2G7^+^ TAMs with a highly selective inhibitor significantly increased CD8+ T-cell activation and efficacy of anti-PD-1 treatment in murine HCC.^54^ Together, these findings highlight possible therapeutic vulnerabilities in PUC warranting further investigation in preclinical models.

The results of this study must be interpreted within the context of the study limitations. First, this study reports on a limited number of PUC tumors, which may limit the power to detect statistical differences and identify rare cell states. Second, the “pure” vs “mixed” classification is based on the tumor area that was evaluated, not the entire tumor. While this distinction is important given the different histologic subtypes included in the sample used and lack of spatial information for the snRNAseq data, a tumor that is 100% PUC histology throughout may show different results than those included in this analysis. Additionally, the spatial neighborhood analysis is based on one tumor, but corroborates the snRNAseq findings. Finally, as in any study performed on human tumor tissue, the findings reported are based on associations and need to be further validated mechanistically. Despite these limitations, this study provides novel insights into PUC biology and tumor microenvironment, biologic insights into possible response to EVP, and identifies novel therapeutic targets for future preclinical testing.

In summary, our work characterizes the TME in PUC and identifies distinct cellular interactions with CDH1-tumor cells and infiltrating immune and stromal cells. We demonstrate that PUC-enriched tumor cells have a luminal transcriptional state, but express cancer stem cell markers, MUC4 and TM4SF1. Additionally, these PUC-enriched tumor cells demonstrate cellular crosstalk with immunosuppressive populations of CAFs (CTHRC1+SFRP2+) and TAMs (TREM2+SPP1+), which may promote a feedback loop that promotes tumor progression and therapeutic resistance. The association between EMT and immune evasion has been shown in other cancers.^56^ Our findings can potentially be utilized to understand PUC response to emerging therapeutics, such as EVP, and identify promising therapeutic vulnerabilities for further exploration.

## Code availability

This study utilized publicly available computing packages. No previously unreported or otherwise custom computer code was generated during this study.

## Acknowledgments

We thank Gabrielle Cannon at the UNC Advanced Analytics Core (Center for GI Biology and Disease; NIH funding number P30 DK034987) for assistance with RNA isolation and library preparation. We thank Yongjuan Xia and Edison Floyd in the Pathology Services Core for expert technical assistance with tissue preparation and Xenium Analyzer spatial omics assay. The PSC is supported in part by an NCI Center Core Support Grant (P30CA016086).

## AI disclosure

The authors used Claude (Anthropic, Sonnet 5.5) to assist with language editing, wordsmithing, and code troubleshooting and editing during manuscript preparation (August through November 2026).

---

**Supplemental Figure 1. snRNAseq identifies general cell types through expression of canonical genes.** Dot plot of select genes stratified by general cell type. Size of the circle indicates percent of cells expressing each gene. Color of circle indicates average gene expression across cells of that subcluster.

**Supplemental Figure 2. Urothelial cell states with low CDH1 expression identified through trajectory inference analysis demonstrate differential expression of genes and molecular pathways.**

(A) Volcano plots depicting quality control metrics (RNA transcript counts, RNA feature counts, and percent mitochondrial) stratified by urothelial cluster.

(B) Volcano plot of 6564 differential expressed between urothelial Cell State 2 vs 3 identified through trajectory inference analysis. Y-axis depicts -log_10_ (adjusted p-value) and x-axis log_2_ fold change.

(C) Bar plot of the differentially expressed Hallmark pathways between Cell state 2 vs 3 identified through trajectory inference analysis. Color of bar indicates p-value.

**Supplemental Figure 3. snRNAseq identifies T-cell subclusters.**

(c) UMAP of T-cell subclusters showing annotated identities.

(D) Dot plot of select genes stratified by T-cell subcluster. Size of the circle indicates percent of cells expressing each gene. Color of circle indicates average gene expression across cells of that subcluster.

(E) Box plots depicting the proportion of T-cells within each tumor comprised by individual T-cell subclusters, stratified by CUC (pink), mixed (green) and PUC (blue) tumors. Boxes indicate the 25^th^ to 75^th^ percentiles, the center line the median, and whiskers extend to 1.5 x IQR; points are individuals tumors. P-values by Wilcoxon rank-sum test.

**Supplemental Figure 4. snRNAseq identifies increase TAM / CD8+ T-cell ratio in PUC tumors vs CUC tumors and distinct macrophage subtypes.**

(A) Boxplot depicting the ratio of TAM to CD8+ T-cells within CUC (left, pink) vs PUC (right, blue) tumors. P-value calculated by Wilcoxon rank sum test.

(B) Boxplot depicting the ratio of CD8+ T-cells to Fibroblasts within CUC (left, pink) vs PUC (right, blue) tumors. P-value calculated by Wilcoxon rank sum test.

(C) UMAP of macrophage subclusters showing annotated identities.

(D) Dot plot of select genes stratified by macrophage subcluster. Size of the circle indicates percent of cells expressing each gene. Color of circle indicates average gene expression across cells of that subcluster.

(E) Box plots depicting the proportion of macrophages within each tumor comprised by individual macrophage subclusters, stratified by CUC (pink), mixed (green) and PUC (blue) tumors. Boxes indicate the 25^th^ to 75^th^ percentiles, the center line the median, and whiskers extend to 1.5 x IQR; points are individuals tumors. P-values by Wilcoxon rank-sum test.

**Supplemental Figure 5. snRNAseq identifies fibroblast subclusters.**

(C) UMAP of fibroblast subclusters showing annotated identities.

(D) Dot plot of select genes stratified by fibroblast subcluster. Size of the circle indicates percent of cells expressing each gene. Color of circle indicates average gene expression across cells of that subcluster.

(E) Box plots depicting the proportion of fibroblasts within each tumor comprised by individual fibroblast subclusters, stratified by CUC (pink), mixed (green) and PUC (blue) tumors. Boxes indicate the 25^th^ to 75^th^ percentiles, the center line the median, and whiskers extend to 1.5 x IQR; points are individuals tumors. P-values by Wilcoxon rank-sum test.

**Supplemental Figure 6. CellChat identifies receptor-ligand pairs predicted to be involved in signaling between fibroblasts, macrophages, and urothelial cells in PUC and CUC tumors.** Bubble plot of ligand-receptors pairs predicted as involved in signaling between the indicated cell types in PUC (pink) vs CUC (blue) tumors. Bubble size indicates p-value. Bubble color indicates communication probability.

**Supplemental Figure 7. Xenium spatial transcriptomics identifies general cell types through expression of canonical genes.** Dot plot of select genes stratified by general cell type. Size of the circle indicates percent of cells expressing each gene. Color of circle indicates average gene expression across cells of that subcluster.

**Supplemental Figure 8. Xenium fibroblast subcluster 2 has a higher expression of immunosuppressive genes also found in PUC-associated fibroblasts in snRNAseq data compared to fibroblasts subcluster 2.** Violin plots show the expression distribution of select genes stratified by fibroblast subcluster in Xenium data. Boxplots within each violin indicate the median (center line), interquartile range (box), and 1.5 × IQR (whiskers); outliers are not shown. P-value calculated by Wilcoxon rank-sum test and is Bonferroni-adjusted.

