## Supplemental Files for "Single Nucleus Profiling Reveals Rewired Tumor-Immune-Stromal Cell Communication in Plasmacytoid Urothelial Carcinoma"

### SUPPLEMENTAL FIGURE 1

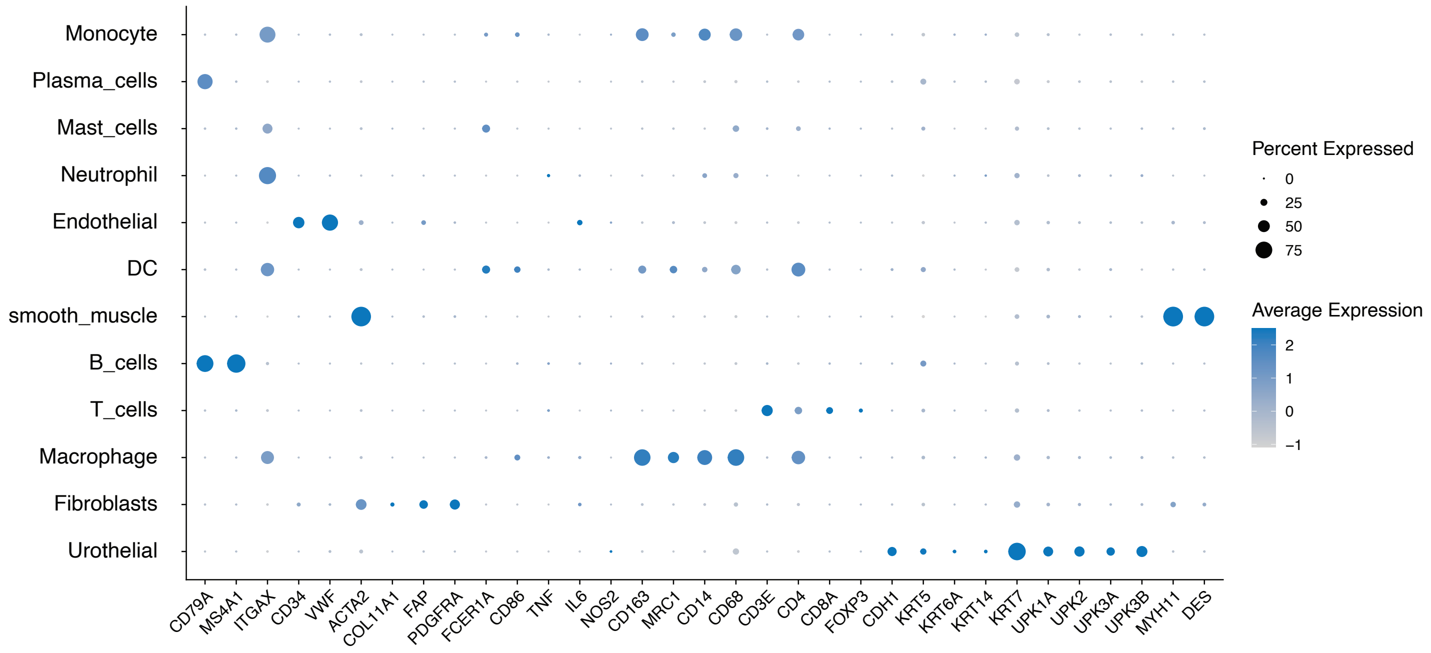

**SUPPLEMENTAL FIGURE 2**

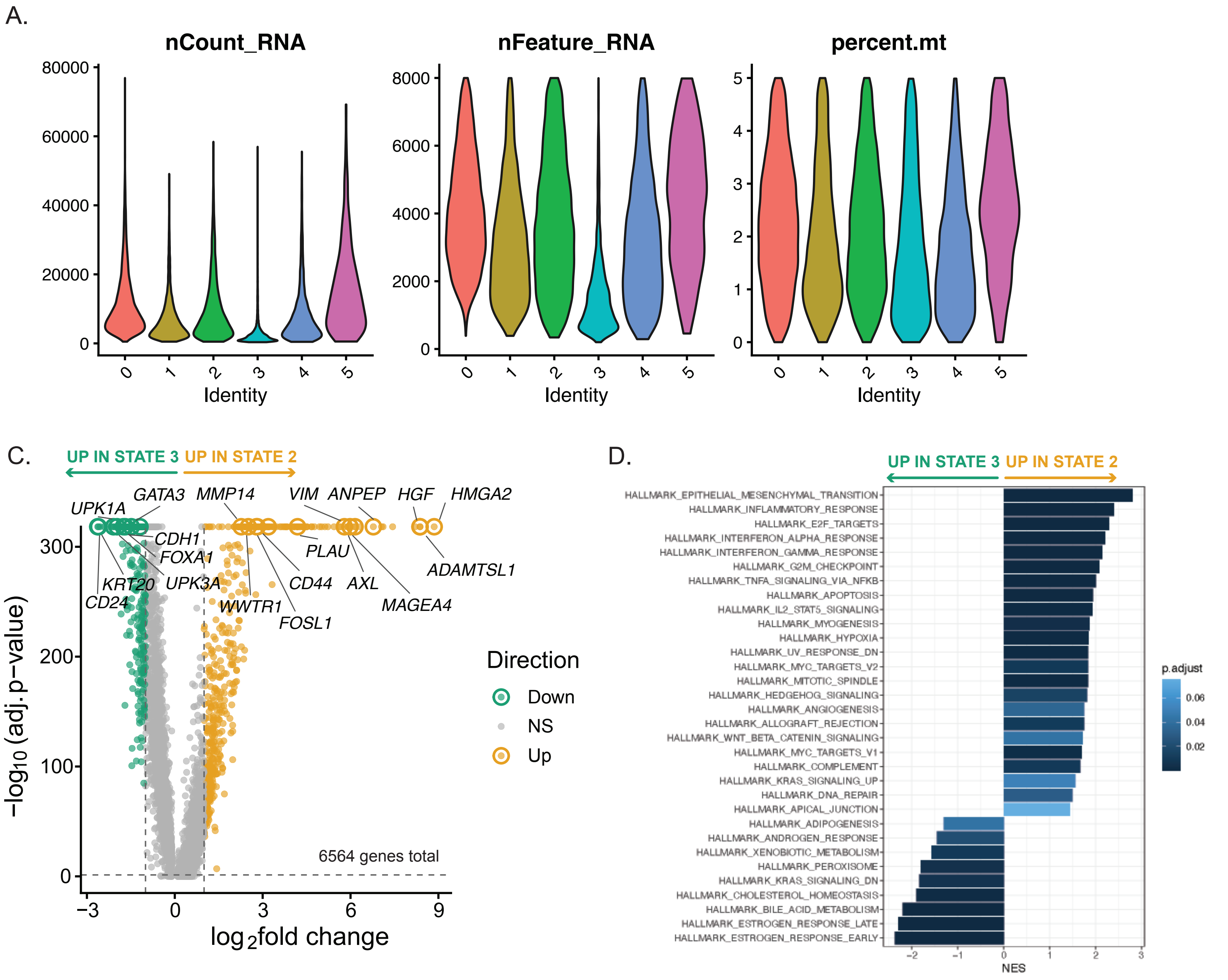

#### SUPPLEMENTAL FIGURE 3

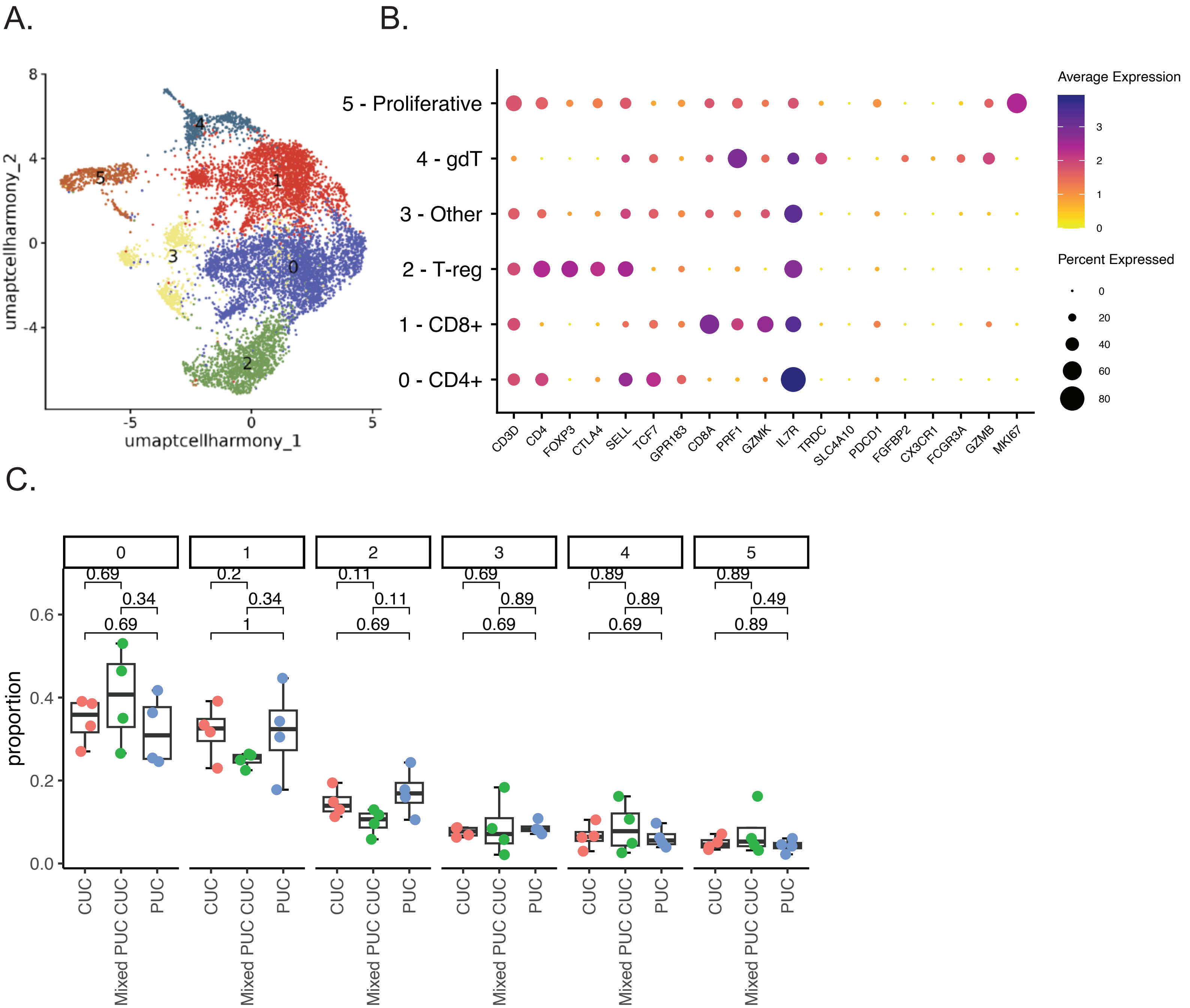

SUPPLEMENTAL FIGURE 4

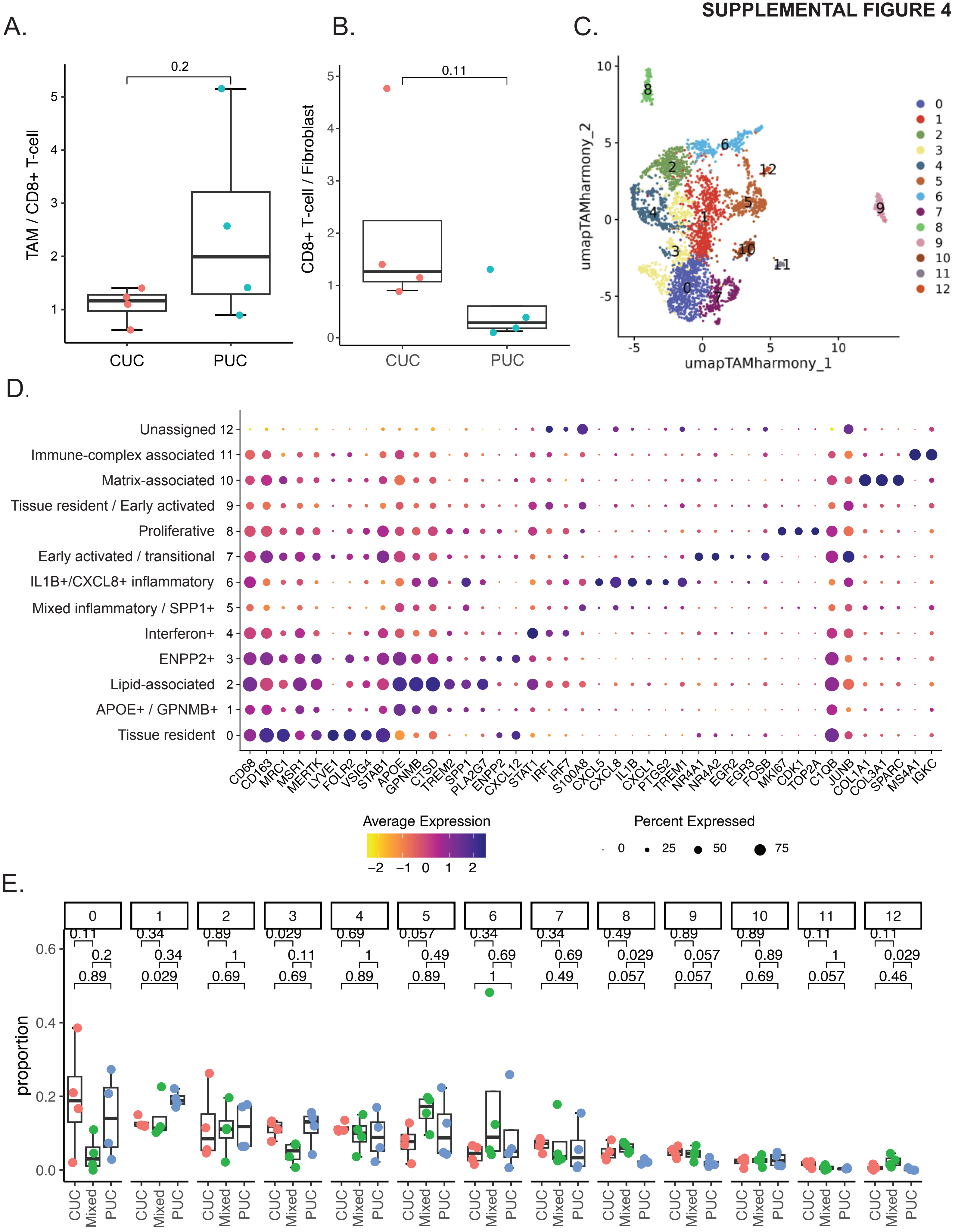

SUPPLEMENTAL FIGURE 5

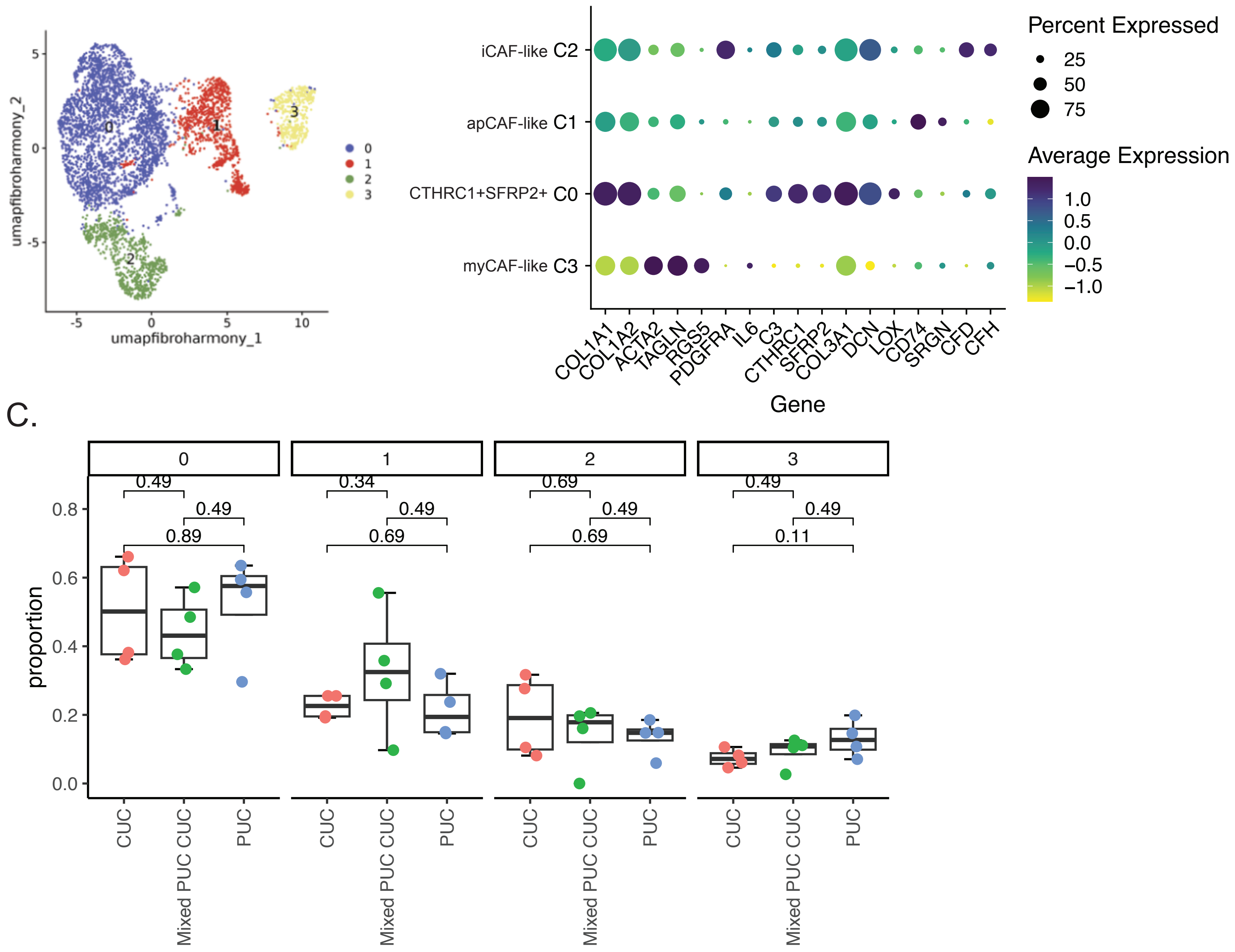

### SUPPLEMENTAL FIGURE 6

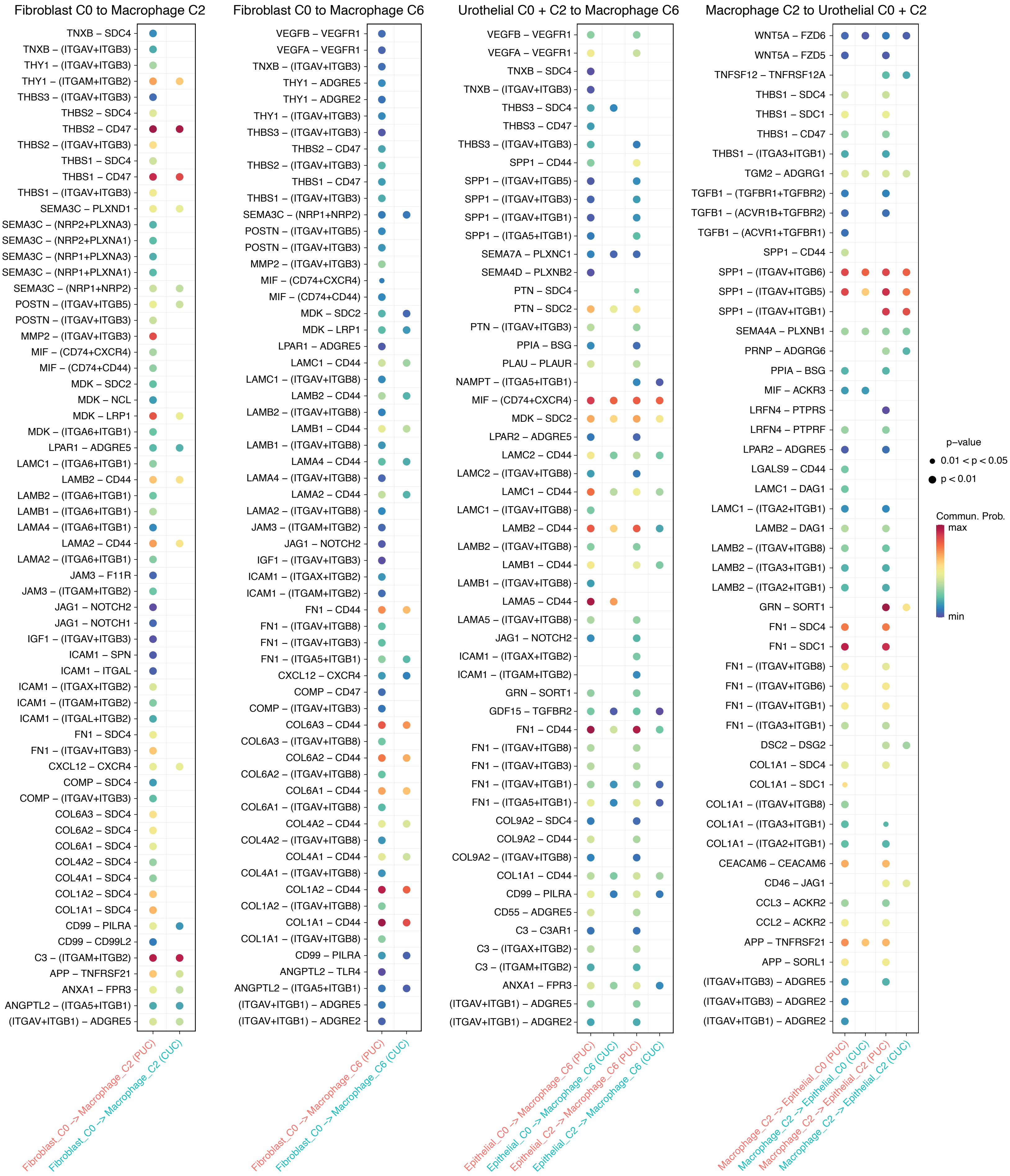

### SUPPLEMENTAL FIGURE 7

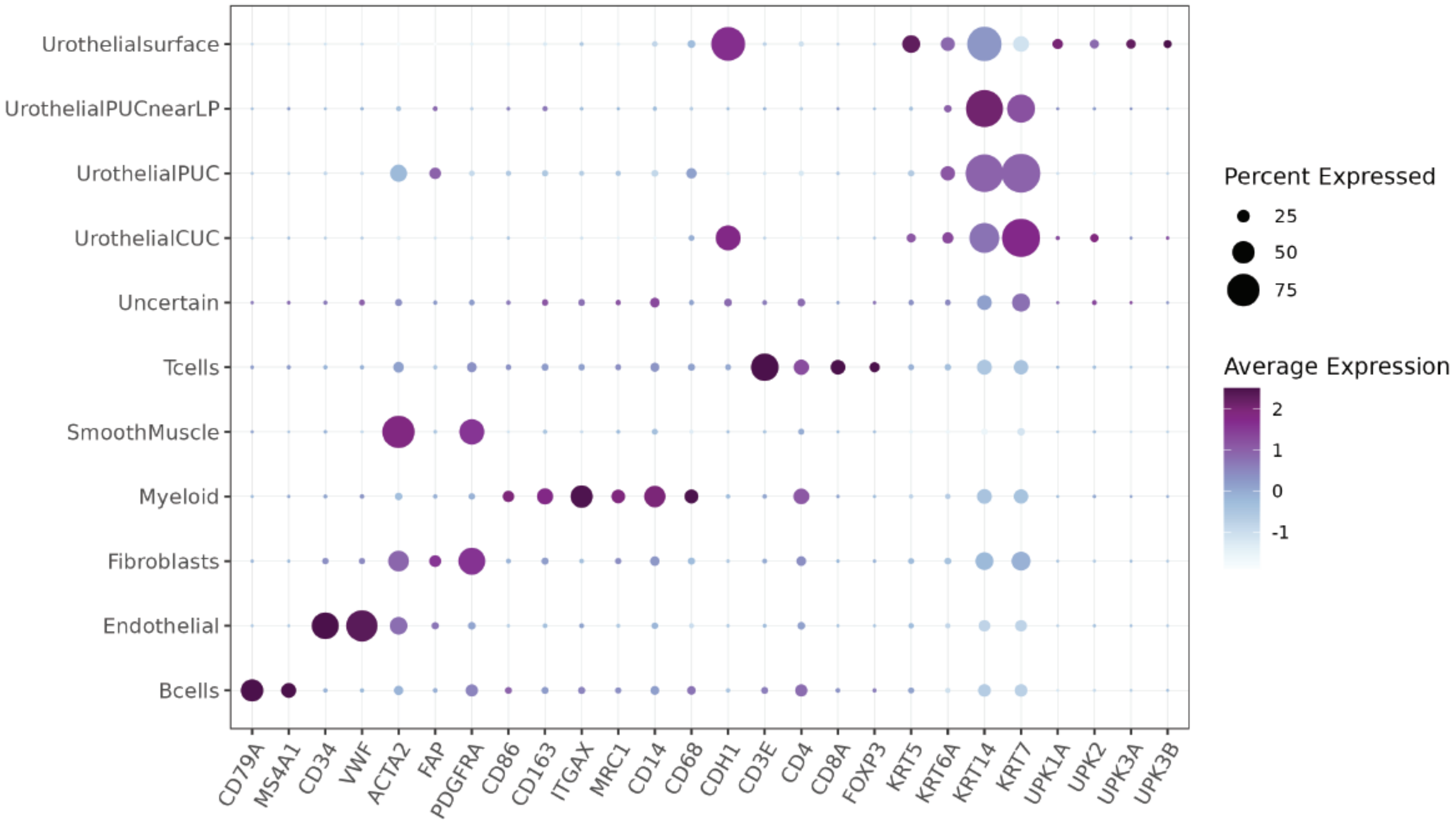

#### SUPPLEMENTAL FIGURE 8

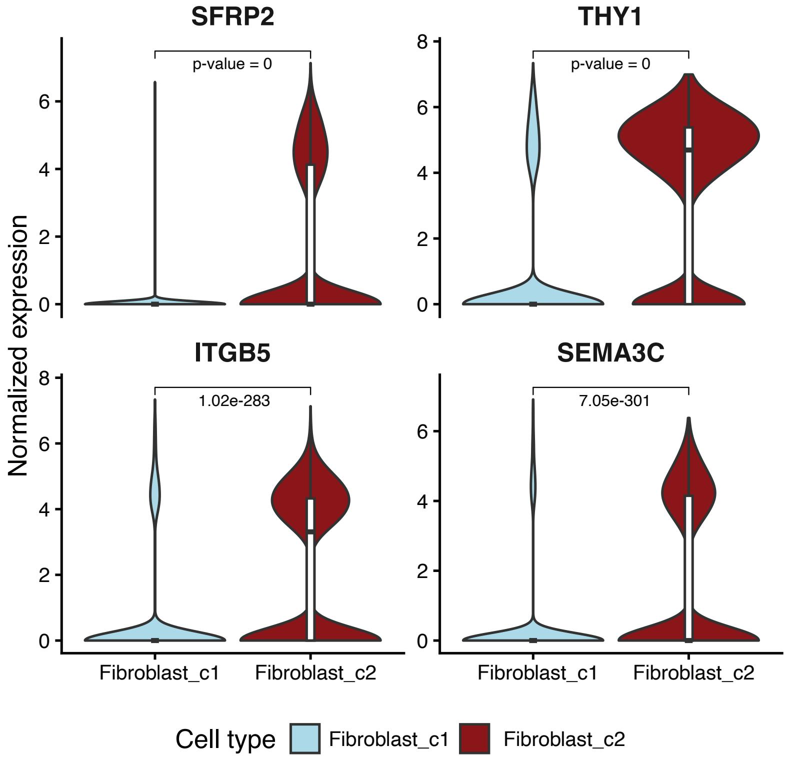
